# Stem Cell Secretome Promotes Scarless Corneal Wound Healing and Rescues Corneal Sensory Nerves

**DOI:** 10.1101/2022.05.07.490347

**Authors:** Ajay Kumar, Yunshu Li, Shayshadri Mallick, Enzhi Yang, Deepinder K. Dhaliwal, Andrew Price, Ting Xie, Yiqin Du

**Affiliations:** Department of Ophthalmology, University of Pittsburgh, Pittsburgh, PA 15213; Stowers Institute of Medical Research, Kansas City, Missouri 64110; Department of Developmental Biology, University of Pittsburgh, Pittsburgh, PA 15213; McGowan Institute for Regenerative Medicine, University of Pittsburgh, Pittsburgh, PA 15213

**Keywords:** Corneal Stromal Stem Cells, Secretome, Wound Healing, Sensory Nerves, Inflammation, Complements

## Abstract

Corneal scarring is a leading cause of blindness in the world. In this study, we explored the therapeutic potential of corneal stromal stem cell (CSSC)-derived secretome in a mechanical debridement mouse model of corneal scarring. CSSC secretome was able to promote scarless corneal wound healing. The mechanisms include 1) dampening inflammation with reduced CD45+, CD11b+/GR1+, and CD11b+/F4/80+ inflammatory cells in the wounded corneas; 2) reducing fibrotic extracellular matrix deposition such as collagen IV, collagen 3A1, SPARC, and α-SMA; 3) rescuing sensory nerves. The proteomic analysis shows upregulated proteins related to wound healing and cell adhesion which boost scarless wound healing. It also shows upregulated neuroprotective proteins in CSSC secretome related to axon guidance, neurogenesis, neuron projection development, and neuron differentiation. Four unique complement inhibitory proteins CD59, vitronectin, SERPING1, and C1QBP found in CSSC secretome contribute to reducing a complement cascade mediating cell death and membrane attacking complex autoantibodies after corneal injury. This study provides novel insights into mechanisms of stem cell secretome induced scarless corneal wound healing and neuroprotection and identifies regenerative proteins in the CSSC secretome.

**Significance Statement:** In this study, we report the therapeutic role of corneal stromal stem cell (CSSC) secretome for scarless corneal wound healing and corneal sensory nerve rescuing. We uncovered that CSSC secretome dampens inflammation, reduces fibrosis, induces sensory nerve regeneration, and rescues corneal cells by inhibiting the complement system in the wounded mouse corneas. This study provides pre-clinical evidence for the use of CSSC secretome as a biologic treatment for corneal scarring to prevent corneal blindness. We delineated a plethora of proteins in the CSSC secretome, which individually or in combination have the potential as future therapies for scarless corneal wound healing.

## Introduction

The cornea is the most anterior part of the eye and provides the most refractive power of the eye. Corneal transparency is vital to vision ^1^. The corneal epithelium, the outermost layer of the cornea, is typically able to heal on its own after damage provided the limbal stem cells are not depleted. When a corneal wound reaches Bowman’s membrane, it ultimately causes damage to the corneal stroma, a mesenchymal tissue making up the thickest layer of the cornea ^2^. Corneal stromal injuries usually cause permanent haze and scarring, therefore leading to vision loss. During corneal wounds, many growth factors and cytokines are activated for wound healing which contribute to keratocytes transformation to fibroblasts and myofibroblasts and excess deposition of extracellular matrix (ECM) proteins and fibrosis ^3^. The cytokines also induce inflammatory and immune cell infiltration into the cornea ^4^. The inflammation also causes corneal cell death and sensory neuron death which compromise corneal transparency and result in vision impairment, even blindness ^5^. Complement cascade is the ultimate outcome of inflammation which causes additional cell death burden on wounded cornea. As of now, the best treatment for corneal scarring with reduced vision is corneal transplantation, which requires donor tissues and includes the risk of graft rejection. Although corneal transplantation is the most successful organ transplantation in our body, long-term success is a challenge as survival rates reduced from 89% at 5 years to 17% at 23 years ^6^. Currently, there are 12.7 million people on the waiting list of corneal transplantation worldwide ^7^. We have shown therapeutic effect of stem cells in the eye in different animal models of eye diseases such as glaucoma ^8,9^ and corneal scarring ^1,10,11^. Corneal stromal stem cells (CSSCs) are located in the corneal limbus close to the corneal limbal epithelial stem cells ^12,13^ and can be separated as a side population ^12^. These cells express stem cell markers CD90, CD73, CD105 and possess ability to differentiate into cells of various lineages ^12,13^. CSSC can suppress corneal scar formation in lumican knockout ^10^ and Col3a1-EGFP ^14^ genetic mouse corneal scar models and prevent corneal scar formation in an acute corneal wound model ^11,15^. This finding opens the door for stem cell-based therapy for the treatment of corneal scars without corneal transplantation. Stem cell secretome, defined as the set of stem cell secreted bioactive factors, including soluble proteins, growth factors, and extracellular vesicles, has regenerative effects ^16^. Currently, sparse studies have investigated the effect of secretome on ocular regeneration and wound healing. In this current study, we investigated the therapeutic effect of CSSC secretome on a mouse corneal wound model and explored possible mechanisms.

## Results

### Corneal stromal stem cell secretome promotes scarless corneal wound healing and dampens inflammation

At first, human CSSC were characterized for their stem cell properties with expression of stem cell markers CD90, CD73, CD105 as per the guidelines of international society of cell therapy ^17,18^ and with additional stem cell markers OCT4, CD166, NOTCH1, STRO1, and ABCG2. CSSC used in the study were >90% expressing CD90, CD73 and CD105, 70-80% expressing ABCG2, NOTCH1, OCT4, STRO1 and CD166, and ∼40% expressing CD271 (*SI Appendix*, Fig. S1 *A-B*). It reflects the stem cell nature of CSSC. Corneal fibroblasts were cultured and characterized with expression of fibronectin and without expression of the stem cell markers as previously described ^8,12^. Secretomes from both CSSC and corneal fibroblasts were collected after they were cultured in serum free conditions for 48 hrs. The cell viability of both CSSC and corneal fibroblasts post secretome harvesting was evaluated by Annexin V/7-Aminoactinomycin D (7-AAD) staining using flow cytometry and live cell stain with viability dyes Calcein Red-Orange and Hoechst 33342. Flow cytometry results showed 3.3±0.7% apoptosis in corneal fibroblasts and 5.1±0.1% apoptosis in CSSC (*SI Appendix*, Fig. S1 *C*) that had no statistical significance. No cell death was observed after Calcein Red-Orange and Hoechst 33342 staining in either corneal fibroblasts or CSSC (*SI Appendix*, Fig. S1 *D*), demonstrating that secretomes were harvested from healthy cells.

Next, it was examined whether the harvested secretome can induce regeneration in wounded mouse corneas. A corneal wound removing the corneal epithelium, Bowman’s member, and superficial stroma was induced using an Algerbrush II as previously described ^11,15,19^. Immediately after wound, 0.5µl of 100U/ml thrombin was applied to the wound site, and1µl of 1:1 mixture of 10mg/ml fibrinogen and 25X concentrated secretome (either from CSSC or fibroblasts) or medium as sham control was added to the thrombin on top of the corneal wound to form fibrin gel attaching to the wounded corneal surface. After a minute, a second round of thrombin and mixed fibrinogen and secretome was applied for enough secretome on the wounded cornea. 72 hrs later, the corneas were examined. Optical coherence tomography (OCT) is a noncontact technology that can be used to produce high-resolution cross-sectional images of cornea located in the anterior segment of the eye ^11^. OCT scanning images showed that the corneas receiving sham treatment and receiving fibroblast secretome treatment showed corneal scar formation suggested by the higher pixels in the images. The CSSC secretome treatment group showed intact epithelial layer and transparent stroma without signs of scar formation comparable to the corneas of naïve control mice (Fig. 1 *A*). The corneas were classified as healed, partially healed, and not healed based on the OCT images. CSSC secretome healed 73.6% corneas (14 out of 19 corneas), partially healed 21% corneas (4 out of 19 corneas), and only 5.2% corneas remained non-healed (1 out of 19 corneas) (Fig. 1 *B*). However, fibroblast secretome was able to heal only 20% corneas fully (4 out of 20 corneas), 52.3 corneas partially (11 out of 20 corneas), and 25% corneas (5 out of 20 corneas) not healing which was comparable to sham controls, healed 15% (3 out of 20 corneas), partially healed 45% (9 out of 20 corneas) and not healed 40% (8 out of 20 corneas).

**Fig. 1.**
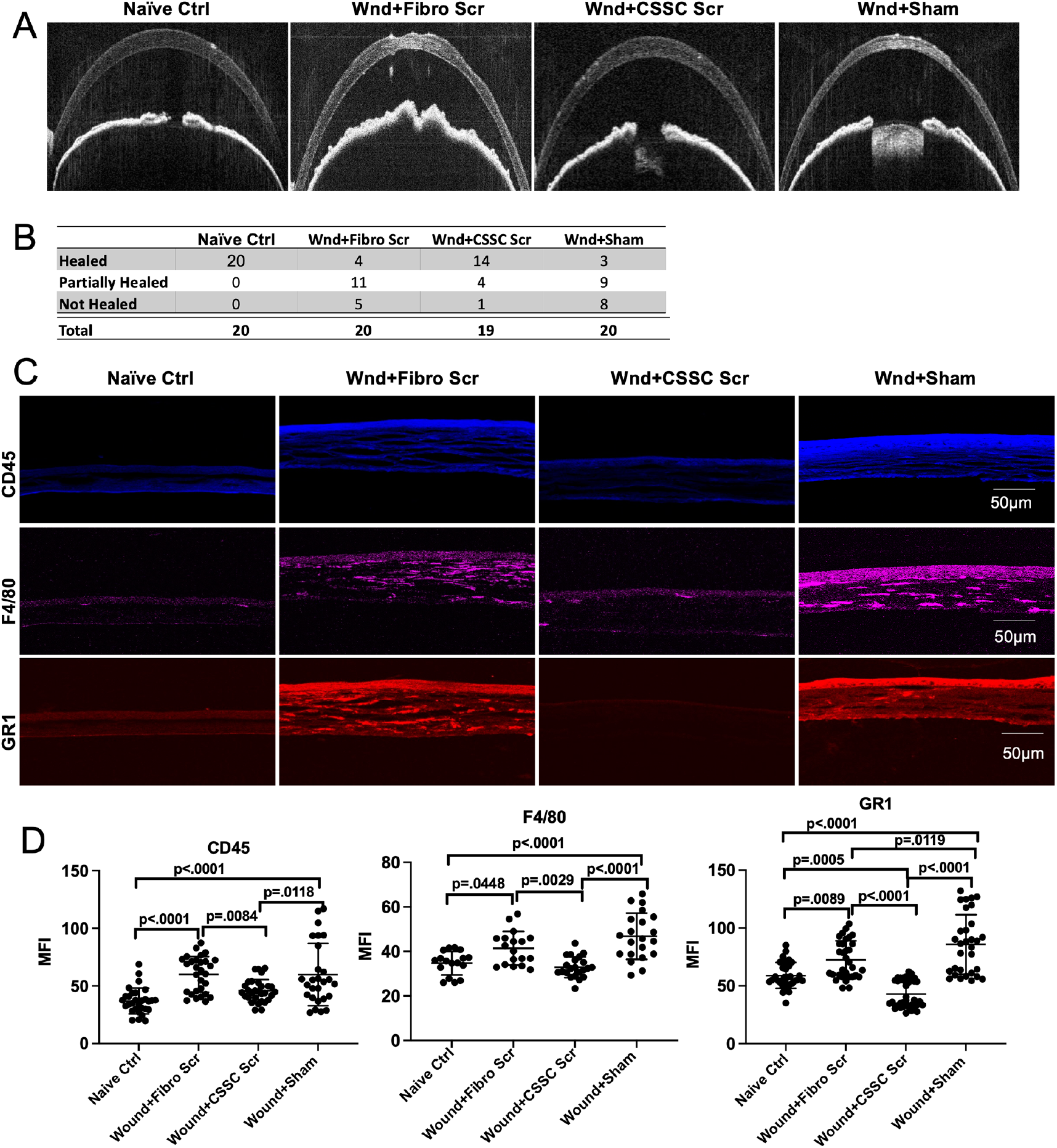
Effect of corneal stromal stem cell secretome on corneal wound healing and inflammation. **A**. Optical coherence tomography (OCT) analysis of mouse corneas showing wounded corneas with different treatments, **B**. Table showing number of corneas with complete healing, partial healing, and no healing (n=19-20), **C**. Immunofluorescent images showing expression of inflammation related antibodies CD45, F4/80, and GR1 in the cornea, scale bars-50µm, **D**. Dot plots represented as Mean±SD for inflammation related antibodies CD45, F4/80, and GR1. Each dot on graph represents mean fluorescence intensity (MFI) from one corneal section. 6-8 corneal sections were imaged per eye (n=3 corneas).

Inflammation plays a critical role in many cellular processes. In vascular tissues it helps to clear pathogen by increasing vascular cell availability to the wounded area, but it can be a liability in an avascular tissue like the cornea and aggravate corneal scarring and damage. The cornea is an immune privileged site, partly due to its avascular nature. Inflammation in the cornea can lead to compromised corneal integrity and promote cell death by infiltration of inflammatory and immune cells, leading to deposition of fibrotic ECM, neovascularization, and tissue destruction ^20^. CSSC have been shown to have anti-inflammatory effects via TSG-6 activation ^21^. Whether CSSC secretome has similar effects to reduce inflammation was examined. CD45 is a commonly used marker for hematopoietic cells except red blood cells and platelets and its increase is associated with activation of inflammatory cells ^1,22^. CD45 expression increased in the mouse corneas of wounded with sham treatment (59.9±27.1) and fibroblast secretome treatment (60.1±15.5) and reduced significantly after CSSC secretome treatment (45.8±9.7), comparable to naïve control corneas (36.9±11.2). This was measured as mean fluorescent intensity (MFI) by immunofluorescent staining (Fig. 1 *C-D*). Immune cell infiltration in the corneas was assessed by immunostaining to detect the expression of F4/80, a well characterized macrophage marker expressed at high level in many macrophages like kupffer cells, microglia etc. CD45^+^/F4/80^+^ cells have been reported to increase after corneal injury ^23^. As shown in Fig. 1 *C-D*, F4/80 expression was significantly increased after corneal wound with sham treatment (46.8±10.4) as compared to naïve control (34.8±5.3) and was significantly reduced after CSSC secretome treatment (32.9±4.7) while fibroblast secretome did not reduce it (41.4±7.5). Another macrophage/myeloid marker Gr1, which stains two Ly6 family Glycosylphosphatidylinositol (GPI) anchor proteins Ly6G (granulocyte marker) and Ly6C (macrophage marker) ^24,25^, was also evaluated. Similarly, Gr1 was significantly increased after corneal wound (91.8±29.6) and reduced after CSSC secretome treatment (42.8±11.7) (Fig. 1 *C-D*). These results indicate that CSSC secretome, not fibroblast secretome, can promote corneal wound healing with dampening inflammation. Analysis of the expression of CD45^+^ (inflammatory cells), CD11b^+^F4/80^+^ (dendritic cells) and CD11b^+^Gr1^+^ (neutrophils) by flow cytometry showed similar results with increased different types of inflammatory cells after corneal wound and with fibroblast secretome treatment, but CSSC secretome reduced CD45^+^ and CD11b^+^F4/80^+^ cell numbers, comparable to naïve control animals (Fig. 2 *A-B*), although CSSC secretome did not reduce the expression of CD11b^+^Gr1^+^ cells. The results indicate that CSSC secretome promotes wound healing and dampens inflammation by preventing infiltration of inflammatory cells on the wounded cornea.

**Fig. 2.**
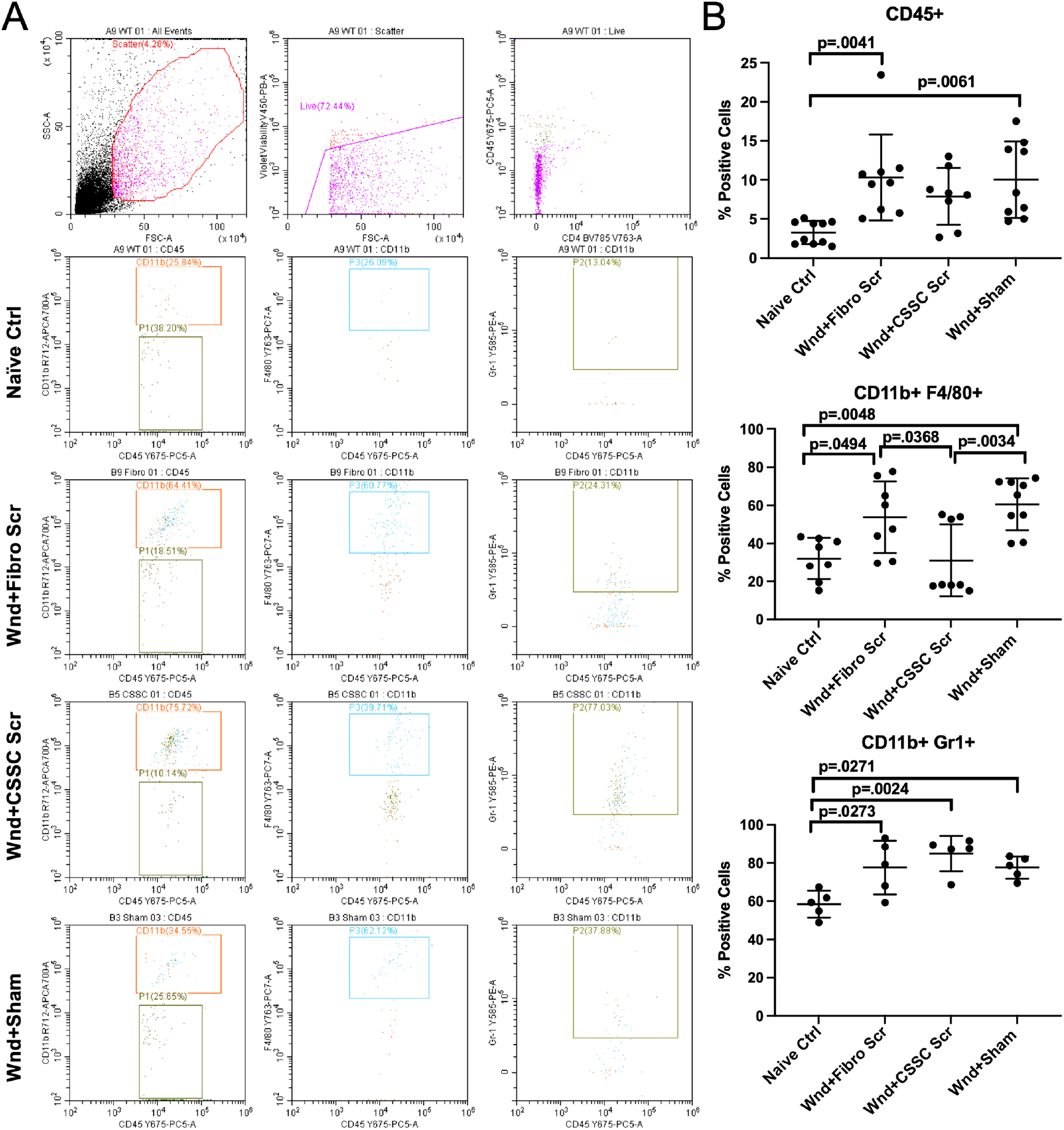
Effect of Corneal stromal stem cell secretome on inflammatory cells. A. Dot plots showing gating strategy for the inflammatory cells in the corneas. Cells were first excluded based on violet live/dead staining. Live CD45+ cells were further gated as CD11b+/- cells and out of those, GR1+/- and F4/80+/- cells were counted, **B**. Dot plots showing percentage positive inflammatory cells for different antibodies, CD45 (n=8-10), CD11b/F4/80 (n=8-9), and CD11b/Gr1 (n=5).

### Stem cell secretome reduces fibrotic extracellular matrix deposition after corneal wound

During corneal wound healing, corneal keratocytes differentiate into fibroblasts and myofibroblasts which secret fibrotic extracellular matrix (ECM) to promote wound healing as well as fibrotic scar formation ^26,27^. To determine whether CSSC secretome alters the ECM deposition and fibrosis during corneal wound healing, the expression of collagen IV (ColIV), collagen 3A1 (Col3A1), and α smooth muscle actin (α-SMA) in the corneas was assessed. ColIV and fibronectin are associated with fibroblastic cells in the injured cornea and corneal scar formation ^28^. ColIV expression was increased by almost 2.5-fold in the scarred corneas treated with sham (26.9±4.8) and with fibroblast secretome (24.8±7.3), as compared to naïve control corneas (10.5±3.6) and was reduced significantly after application of CSSC secretome (20.1±2.6) (Fig. 3 *A-B*). Col3A1 is highly expressed in scarred cornea and is widely used as a marker of corneal stromal fibrosis ^14^. Similarly, CSSC secretome reduced the Col3A1 expression (22.5±5.4) which was increased in the wounded corneas with sham treatment (30.6±5.1) and with fibroblast secretome treatment (29.2±5.9) (Fig. 3 *A-B*). Secreted protein acidic and rich in cysteine (SPARC) is a collagen binding protein and binds to collagen III and IV by its E-C domain to facilitate their assembly to ECM ^29,30^. SPARC expression is increased during corneal fibrosis as shown previously ^18,27,31^. SPARC expression was significant increased in the wounded corneas treated with sham (37.6±9.3) and with fibroblast secretome (33.8±5.5), as compared to naïve control corneas (24.4±7.3). CSSC secretome treatment reduced SPARC expression significantly (25.2±8.0) (Fig. 3 *C-D)*. α-SMA is expressed by activated fibroblasts and myofibroblasts in wounded corneas ^32^. The expression of α-SMA was almost doubled after corneal wound with sham treatment (34.0±11.3) as compared to control (18.6±4.1) (Fig. 3 *C-D*). Interestingly, fibroblast secretome worsened the condition by increasing the expression of α-SMA almost three-fold (51.0±16.6). CSSC secretome (30.9±7.5) did not induce any significant reduction of α-SMA as compared to the sham control. This finding emphasizes that CSSC secretome does induce regeneration in wounded corneas by reducing the expression of fibrotic markers and fibrotic ECM deposition. A comparative label-free protein identification and quantification (proteomics) by liquid chromatography coupled tandem mass spectrometry (LC-MS/MS) was performed on the secretomes from CSSC and corneal fibroblasts. Interestingly, CSSC secretome contains 41 proteins that promote corneal wound healing and fibroblast secretome only contains such 18 proteins related to corneal wound healing (*SI Appendix*, Fig. S2 *A*). CSSC secretome also contains a high number of cell-cell adhesion proteins or molecules while fibroblast secretome contains fewer such proteins (*SI Appendix*, Fig. S2 *B*), indicating these cell-cell adhesion proteins may contribute to the scarless wounded healing promoted by CSSC secretome.

**Fig. 3.**
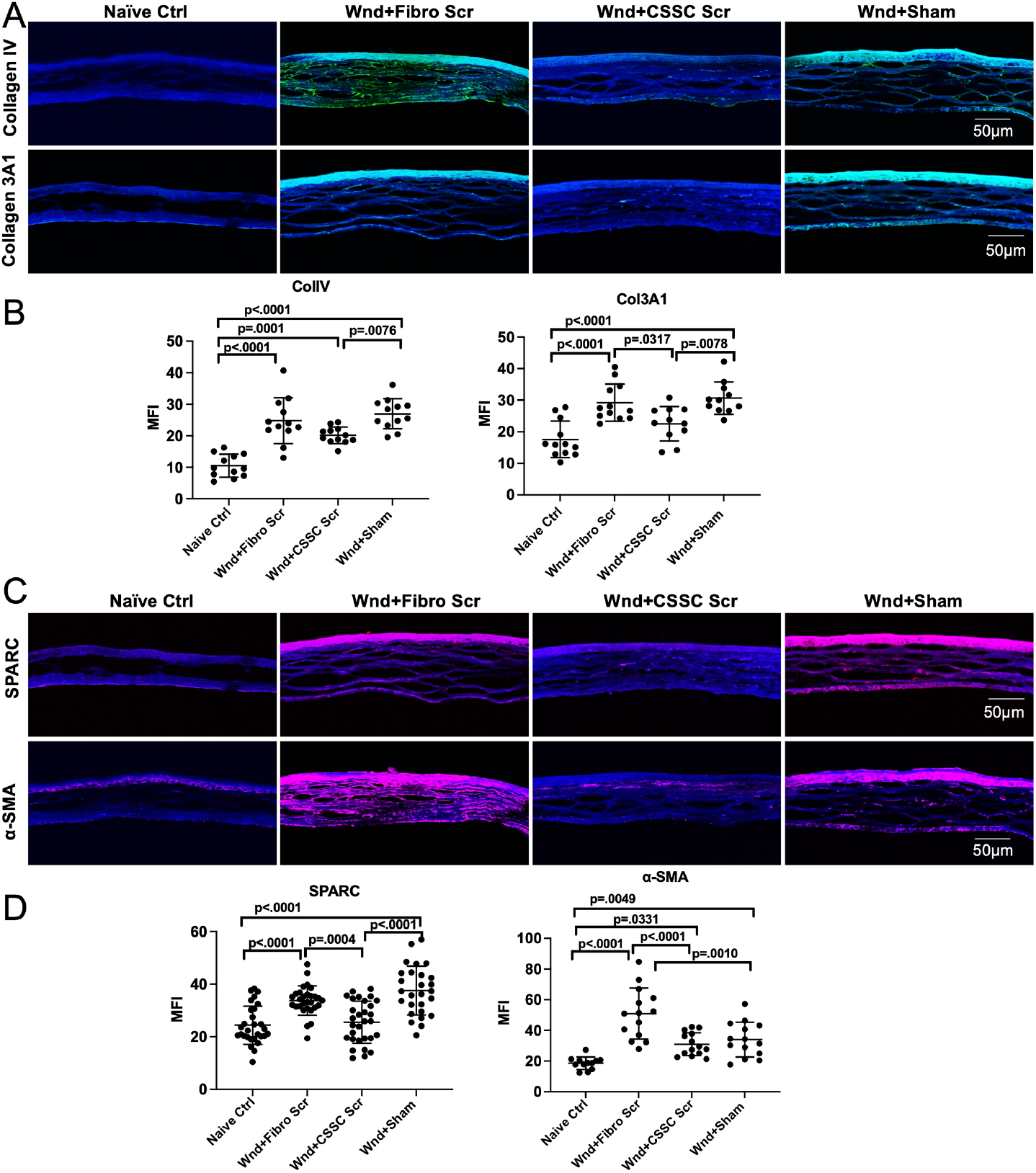
Effect of corneal stromal stem cell secretome on extracellular matrix (ECM) deposition and fibrosis. **A-B**. Immunofluorescent images and dot plots showing expression and quantification of mean fluorescence intensity (MFI) for ECM markers Col IV and Col3A1 in the cornea, **C-D**. Immunofluorescence images and quantification of MFI for the fibrotic markers SPARC and α-SMA in the cornea, scale bars-50µm. Dot plots are represented as Mean±SD. Each dot on graph represents MFI from one corneal section. 6-8 sections were photographed per eye (n=3 corneas).

### Stem cell secretome promotes corneal sensory nerve regeneration

One of the major complications resulting in vision loss in corneal wound is loss of sensory neurons. Infiltration of immune cells in the cornea usually causes death of sensory neurons ^1,33^. Blocking this interaction might prevent corneal opacity ^5^. Death of sensory neurons may lead to loss of blinking response which can inhibit tear production and distribution. This can result in dry eye due to corneal desiccation and aggravation of corneal inflammation and scarring. Since CSSC secretome can prevent inflammatory cell infiltration and dampen inflammation, it might be able to protect sensory nerves. Two neuronal markers β-3 tubulin and P-substance were examined in the whole mount corneas by immunostaining. Overall, naïve control corneas showed plenty of sensory nerve fibers in the cornea with positive β-3 tubulin staining (106.1±23.6, Fig. 4 *A*). The wounded corneas that received CSSC secretome had partially regenerated sensory nerve plexus (76.2±21.4), whereas fibroblast secretome (44.2±17.8) and sham treatment groups (51.1±11.4) showed less sensory nerve fibers (Fig. 4 *A*). Similarly, the expression of P-substance, a sensory neuron marker, was significantly increased in the wounded mouse corneas that received CSSC secretome treatment (97.06±19.6) in comparison to that treated with sham (62.99±14.4) and with fibroblast secretome (54.81±16.3), which were reduced as compared to the naïve control (74.61±11.41) (Fig. 4 *B*).

**Fig. 4.**
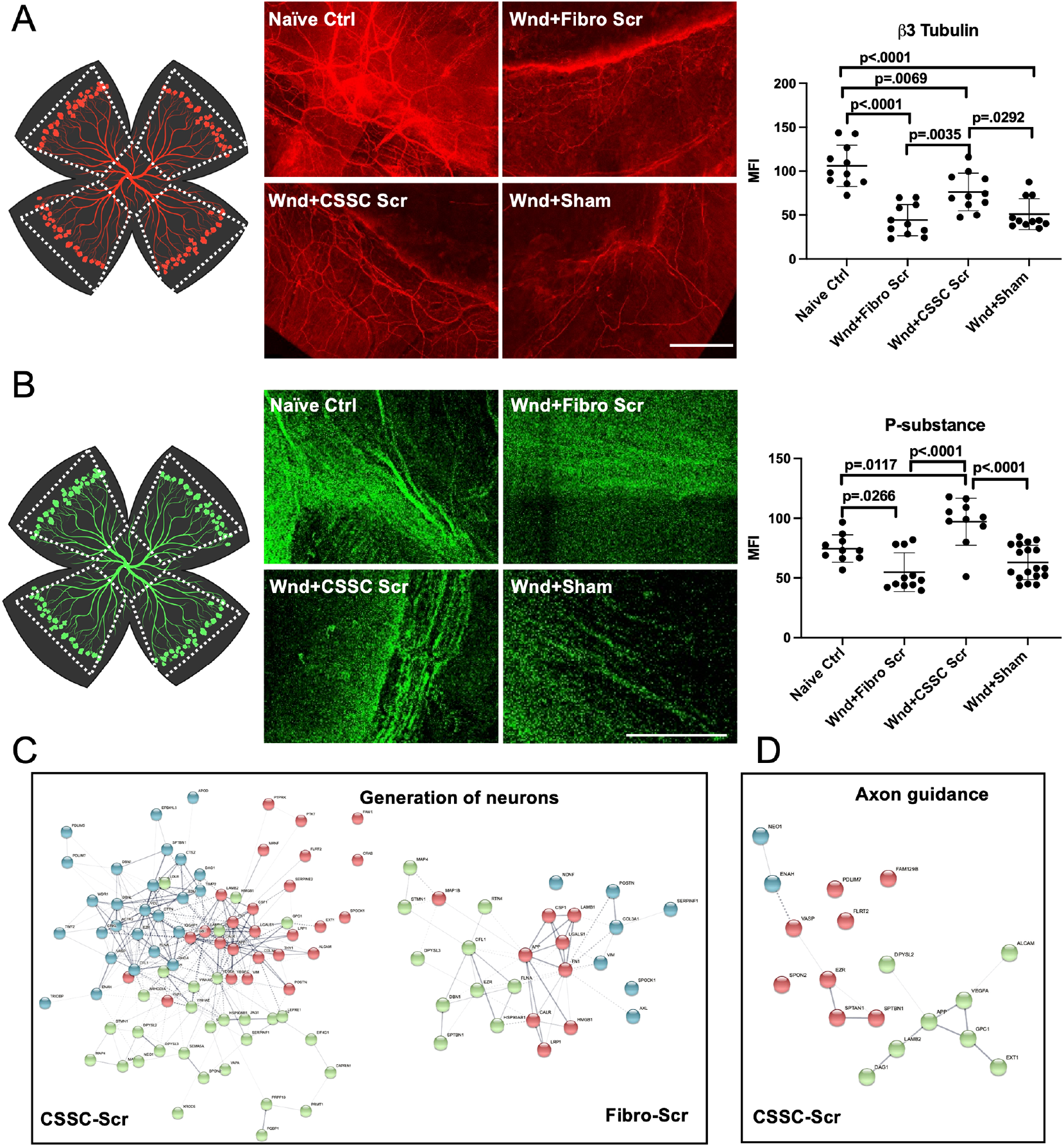
Corneal stromal stem cell secretome induce sensory nerve regeneration by key proteins. **A**. A schematic representing wholemount cornea and the scheme for how corneal neurons were quantified for the mean fluorescence intensity (MFI) of β-3 tubulin (image adapted from Birender.com, left panel). Immunofluorescent stitched images composed of multiple z-stacks acquired for the expression of neuronal marker β-3 tubulin in wholemount corneas (middle panel). Corneal nerve plexuses are clearly visible in naïve control and CSSC secretome treated corneas whereas dramatically lost in sham. Dot plots (right panel) showing quantification of MFI of β-3 tubulin staining. **B**. Schematic represent wholemount cornea and how MFI was quantified for p-substance (image adapted from Birender.com, left panel), confocal images showing stitched z-stacks for expression of P-substance in wholemount corneas (middle panel), Dot plots (right panel) showing quantification of MFI of P-substance staining. **A-B**. One dot on the bar graphs represents quantification from one ribbon of the cornea (n=3 corneas). Data is represented as Mean±SD. **C-D**. Interactome analysis of secretome proteins from both CSSC and fibroblasts, showing interaction between proteins involved in generation of neurons in both CSSC and fibroblasts and axon guidance in CSSC, as analyzed by String v11, (n=2 each).

Proteomic data show that 82 different proteins in CSSC secretome pertain generation of neurons Proteomic data show that 82 different proteins in CSSC secretome pertain generation of neurons such as TIMP1, TIMP2, SEMA5A, while in fibroblast secretome, only 26 of such proteins were identified (Fig. 4 *C*). Axon guidance proteins guide axons during development so that neurons can navigate to their specific targets in a complex microenvironment. 18 axon guidance proteins were found in CSSC secretome but none in fibroblast secretome (Fig. 4 *D*). Further proteomic analysis showed more neural differentiation proteins and proteins involved in the development of neuron projections as compared to fibroblast secretome (*SI Appendix*, Fig. S3 *A-B*). Cumulatively, the proteins involved in wound healing, generation of neurons, axon guidance, neuron projection development and neuron differentiation might enhance the overall therapeutic effect of CSSC secretome on corneal wound healing and neuron regeneration (*SI Appendix*, Fig. S3 *C*).

### Stem cell secretome rescues corneal cell death by inhibiting complement system

Corneal wounds induce a massive cell death in the cornea by promoting inflammation and apoptosis ^3^. To detect if CSSC secretome can protect corneal cells from death in wounded corneas, terminal deoxynucleotidyl transferase dUTP nick end labeling (TUNEL) staining was performed on the cryosectioned corneal tissue to detect DNA fragmentation by labeling the 3′-hydroxyl termini in the double-strand DNA breaks in apoptotic cells. There was approximately 15-fold higher cell death in wounded corneas treated with sham (146.0±25.9) as compared to naïve control corneas (10.3±4.2) (*SI Appendix*, Fig. S4 *A-B*). CSSC secretome treatment (98.9±20.3) significantly reduced the number of TUNEL+ cells undergoing apoptosis, while fibroblast secretome did not reduce the TUNEL+ cell number significantly (124.5±29.7). Since wounded corneas have more inflammatory cells and generally, inflammation is mediated by activated complements, it was questioned whether there is any involvement of complements in corneal wound healing by CSSC secretome. The proteomic analysis revealed a remarkable difference in the complement related proteins in CSSC and fibroblast secretomes (Fig. 5 *A*). CSSC secretome expressed four unique complement proteins CD59, SERPING1, C1QBP, and vitronectin while fibroblast secretome had no expression on any of these proteins (Fig. 5 *A*). Surprisingly, all these four proteins inhibit complements. Complements usually causes formation of autoantibodies, leading to cell death. We detected autoantibody accumulation on cells by staining with anti-mouse IgG antibody. Naïve control corneas showed minimal autoantibody formation on the corneal cells (17.3±9.3) which was significantly increased in wounded corneas treated with sham (31.2±14.5, p=0.0151). CSSC secretome treatment reduced the level of autoantibodies (29.0±10.9) comparable to the naïve control (p=0.0521). Fibroblast secretome treatment dramatically increased the autoantibody level (51.7±17.7, p<0.0001) (Fig. 5 *B*). This led us to investigate the involvement of proteins regulating complements in the secretomes. Two such proteins CD59 and VTN in the mouse corneas were examined by immunostaining. CD59 expression was significantly reduced in wounded corneas treated with sham (20.8±3.9) and with fibroblast secretome (20.2±6.9) as compared to naive control (30.4±12.9), and was significantly increased (27.8±5.2) after CSSC secretome treatment, comparable to the level in naïve control (p=0.7205) (Fig. 5 *C-D*). Interestingly, vitronectin expression was not altered significantly in wounded corneas after treatment with sham (35.9±6.2) or fibroblast secretome (41.6±13.4) as compared to naive control (40.2±9.6). CSSC secretome treatment on the wounded corneas significantly increased the vitronectin expression (45.2±8.4) as compared to sham treatment (p=0.0303, Fig. 5 C-*D*). The increased expression of CD59 and vitronectin may provide additional protection to corneal cells from complement-induced cell death.

**Fig. 5.**
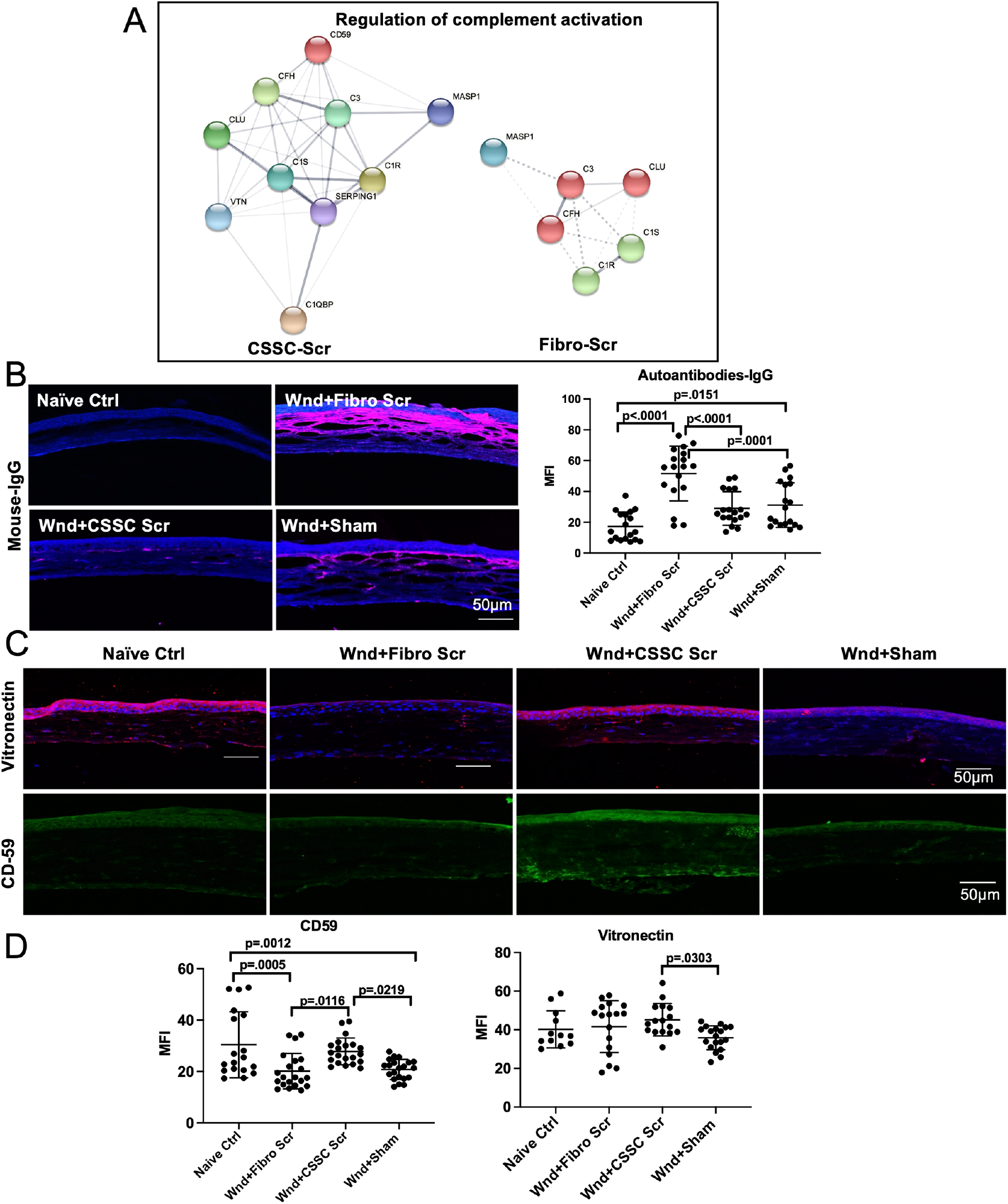
Corneal stromal stem cell secretome inhibits complements. **A**. Interactome analysis for secretome proteins from both CSSC and fibroblasts showing presence of four unique proteins CD59, Vitronectin (VTN), SERPING1, and C1QBP in the CSSC secretome as analyzed by String v11, (n=2 each), **B**. Immunofluorescent images showing autoantibody staining using a goat anti-mouse IgG antibody and dot plots showing mean fluorescence intensity (MFI) quantification showing a higher formation of autoantibodies in sham and fibroblast secretome treated corneas, **C**. Immunofluorescent images showing increased expression of vitronectin and CD59 in the control and CSSC secretome treated corneas while sham and fibroblast secretome treated corneas displayed reduced expression, **D**. Dot plots represent quantification of the MFI for CD59 and vitronectin. Scale bars-50µm. Data is represented as Mean±SD. Dots on graph represent MFI from one corneal section. 6-8 sections were photographed per eye (n=3).

In nutshell, our results showed that CSSC secretome induces corneal wound healing by dampening inflammation, reducing fibrotic ECM deposition, enhancing sensory neuron regeneration, and preventing cell death by proteins involved in cell-cell adhesion, wound healing, neuroprotection, and complement inhibition (Fig. 6).

**Fig. 6:**
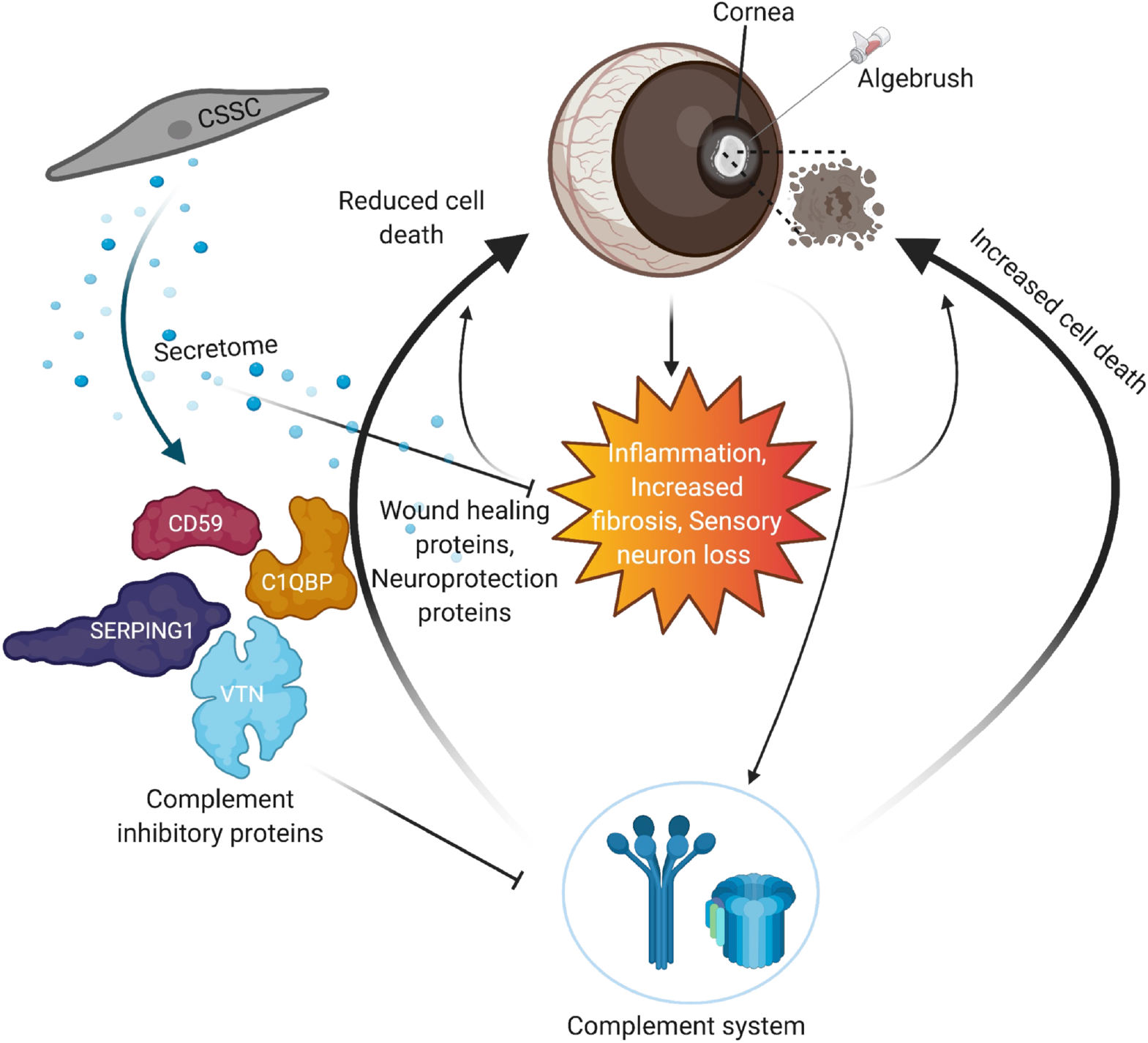
Proposed model of corneal regeneration by CSSC secretome. Induction of corneal wound increases inflammation and fibrosis which result into increased cell death in the wounded corneas. Corneal wound also increases death of sensory neurons in the cornea which enhances corneal opacity. Neuroprotective proteins present in CSSC secretome related to neurogenesis, neuron projection development, and neuron differentiation rescue the neuronal loss in the wounded corneas. Also, proteins related to wound healing in CSSC secretome provide support for scarless wound healing. Complement inhibitory proteins present in CSSC secretome like CD59, SERPING1, C1QBP, and vitronectin (VTN) inhibit the complement system, rescuing the corneal cell death, and promoting scarless corneal wound healing.

## Discussion

In this study, we used a mouse corneal wound model to exhibit the therapeutic effect of CSSC secretome on promoting scarless wound healing and increasing sensory nerve regeneration. Inhibition of complement systems plays a role in scarless wound healing. Proteomic data indicate some important proteins in the CSSC secretome involved in the process, such as CD59, C1QBP, vitronectin, SERPING1 in promoting corneal wound hearing; TIMP1, TIMP2, SEMA5A in promoting neuronal regeneration; NEO1 and APP in neuron projection development and axon guidance.

Similar to our previously reports ^1,18^, the CSSC used in this study expressed the stem cell markers CD90, CD73, CD105, OCT4, ABCG2 etc., indicating their stemness. Our group and others have reported the potential of CSSC to promote corneal wound healing in a variety of corneal scarring models ^10,11,15,34^. Previous reports have explored the role of stem cell-derived paracrine factors and extracellular vesicles (EVs) for corneal wound healing by upregulation of Akt signaling pathway ^35^ or by delivery of miRNAs ^36^. There have been sparse studies reporting stem cell secretome which spans both EVs and soluble proteins present in cells’ secretion cargo. Our report on the ability of CSSC secretome to promote scarless corneal wound healing successfully suggests that secretome alone can recapitulate the wound healing effect of CSSC and thus, maybe a better approach for corneal wound healing due to non-involvement of cellular part. Also, CSSC secretome contains multiple proteins involved in corneal wound healing and cell adhesion, such as CD59, C1QBP, vitronectin, and SERPING1. During corneal wound, corneal epithelial cells disassemble their hemidesmosomes and migrate as a sheet to cover the wound by activating focal contact, a type of adherent junction ^37^. This junction is present only at the end of migrating cells. The upregulation of key cell adhesion proteins in CSSC secretome as compared to fibroblast secretome might be involved in increasing the adherent junctions in the migrating corneal cells and support more effective wound healing.

Inflammation is the body’s active and reactive defense mechanism which stems from an efficient non-specific process of its response to a host of pathogens. This process mainly involves infiltration of white blood cells and other immune cells including macrophages to the injured site to clear pathogens which results in symptoms such as redness, warmth, and swelling. Inflammation plays a context dependent role; it is beneficial in vascular tissues but harmful in avascular tissues like cornea. In response to inflammation, the cornea can develop different pathologies like superficial punctate keratitis, epithelial erosions, corneal swelling, neovascularization etc., if left untreated ^20,38^. We have reported previously that after corneal wound there is increased infiltration of CD11b^+^/Ly6G^+^ neutrophils in corneas at 24 hr which was reduced in CSSC-treated wounds ^21^. Also, CD45^+^/CD11b^+^/Ly6C^+^ monocytes have been reported to increase just after 6 hrs. of dulled blade wound induction in the cornea ^23^. The ability of CSSC secretome to reduce the number of CD45^+^/CD11b^+^Gr^+^/CD11b^+^F4/80^+^ cells indicate that CSSC secretome may be an ideal candidate to reduce corneal inflammation and clear corneal opacity. One of the major complications of corneal wound healing is fibrosis which is mainly mediated by transforming growth factor (TGF-β), which is involved in myofibroblast conversion and fibrotic ECM formation ^3,39^. Although there are a few anti-TGF-β drugs approved for human use like ROCK inhibitors ^40^, Rosaglitazone ^41^, and Trichostatin A ^42^, we are still far away from finding a biologic treatment which can become part of routine clinical practice to prevent fibrotic ECM and provide scarless wound healing. We showed recently that CSSC derived secretome can reduce fibrosis markers SPARC, CTGF, and fibronectin and increase corneal wound healing in cultured human corneal fibroblasts in vitro ^18^. The ability of CSSC secretome to effectively reduce both inflammation and fibrosis in the current corneal wound model provides a novel treatment strategy to use CSSC secretome for effective scarless corneal wound healing.

Inflammatory cells have been reported to induce corneal opacity and application of steroids or use of surgical techniques is often employed to reduce these cells and treat corneal opacity ^43^. However, these treatments are unable to regenerate sensory neurons and complete clearance of immune cells might not be possible if there is no sensory nerve regeneration. Yun et al. showed that sensory nerve regeneration can be achieved in wounded corneas by removing sympathetic nerve using superior cervical ganglionectomy, which can restore corneal transparency ^43^. That study showed that regeneration of sensory nerves at the last stage of corneal disease, when corneal transplantation is the only option, can rescue corneal transparency. The infiltration of immune cells in the scarred cornea can produce various pathogenic cytokines like vascular endothelial growth factor-A (VEGF-A) which might induce neovascularization and innervation of sympathetic nerve fibers in cornea which increase corneal opacity ^5^. CSSC secretome effectively reduces the infiltration of inflammatory and immune cells in the wounded corneas which results into regeneration of sensory nerves, providing an effective alternative to complex surgical techniques and an easy approach to restore corneal transparency. A detailed investigation of CSSC and fibroblast secretome provided us novel insight into CSSC secretome mediated neuroprotection. Plethora of proteins present in CSSC secretome providing axon guidance cues, proteins involved in neuron projection development and differentiation, might further explain the therapeutic role of CSSC secretome in corneal wound healing. Interestingly CSSC secretome expressed 18 different axon guidance proteins whereas fibroblast secretome expressed none. These axon guidance proteins provide neuroprotection by guiding neurons to their specific targets. e.g., neogenin (NEO-1) found in CSSC secretome interacts with netrin-1 and acts as an axon guidance molecule in vivo and the neurons expressing NEO-1 can navigate their ventral trajectory by using several attractive and repulsive cues ^44^. Another unique protein in CSSC secretome, amyloid precursor protein (APP) is involved in both neuron projection development and axon guidance. Although APP’s role has been focused mainly on the pathogenesis of Alzheimer’s disease, recent trauma research has shown the considerable neuroprotective role of APP in traumatic brain injury models ^45^, emphasizing it as an ideal future therapeutic candidate for neuroprotection. Collectively, CSSC secretome showed presence of hundreds of these neuroprotective proteins which might be used as effective small molecule based therapeutic modalities individually or in combination for neuroprotection and in regenerating sensory neurons in wounded corneas to restore vision and in other optic neuropathies like glaucoma and other neurodegenerative disorders.

One of the reasons for secretome mediated neuroprotection might be inhibition of complements by proteins present in CSSC secretome. Complement inhibition has been shown to be neuroprotective after cerebral ischemia and reperfusion ^46^. Complement CD59 gene encodes a ubiquitously expressed membrane-bound glycoprotein which inhibits complement system. Mechanistically, CD59, also called membrane attack complex (MAC) inhibitory protein, binds complement C8 and/or C9 during MAC formation, thus preventing cells from generating autoantibodies, which cause cell death, kill themselves, and inhibiting the terminal pathway of complement cascade ^47,48^. Reduction of CD59 after corneal wound and its increase after CSSC secretome treatment potentially reflects a novel role for CSSC secretome for protection of cell death and promotion of corneal wound healing. A recent study reported that mesenchymal stem cell exosomes could inhibit complement activation and break the feed-forward loop of complements and neutrophils via exosome CD59 in severe COVID-19 patients ^49^. This finding is similar to what we report here. Vitronectin is found in serum and tissues and promotes cell adhesion and spreading, inhibits the membrane-damaging effect of the terminal cytolytic complement pathway, and binds to several serpin serine protease inhibitors. SERPING1 gene encodes a highly glycosylated plasma protein involved in the regulation of the complement cascade. SERPING1 encoded protein, C1 inhibitor, inhibits activation of C1r and C1s, which are the first complement components, and thus inhibits complements ^50^. During complement activation, C1r and C1s are associated with another complement component C1q to form the first component of the serum complement system. C1QBP gene encoded protein further inhibits C1 activation by binding to the globular heads of C1q molecules. Normally the cornea maintains very high levels of complement inhibitory proteins CD59, decay accelerating factor (DAF, CD55), and membrane cofactor protein (MCP, CD46), all strongly expressed in the corneal epithelium and also to somewhat extent in corneal stroma ^51^. Loss of these proteins after corneal wound might lead to activation of complements and increased cell death. CD59 is a cell surface protein and is attached to cell membrane by a glycosylphosphatidylinositol (GPI) tail. CD59 is the only membrane-bound inhibitor of the terminal pathway of complement cascade and prevents self-destruction of cells. Recently, complement modulation has been shown to reverse the pathology of retinal pigment epithelial cells in age-related macular degeneration ^52^. Our study suggests that complement inhibition might be equally beneficial in promoting wound healing and is one of the important mechanisms by which CSSC secretome induces corneal wound healing and neuroprotection. We recently reported the role of a different stem cell type, trabecular meshwork stem cell secretome (TMSC) in the protection of retinal ganglion cells in ocular hypertension glaucoma mouse models ^53^. Since CSSC secretome also harbors these neuroprotective proteins, it might be equally effective for the treatment of other optic neuropathies like glaucoma.

In nutshell, this study reports advancement over previous approaches towards corneal wound healing and provides insight into the novel mechanisms of scarless corneal wound healing by stem cell secretome. Further large animal studies and clinical trials would warrant the safety and efficacy of this approach in human.

## Materials and Methods

### Cell culture and collection of secretome

Human CSSCs were obtained from donor corneas and cultured in Dulbecco’s modified Eagle’s medium (DMEM) low glucose mixed with different growth supplements as reported before ^12,18^. Human corneal fibroblasts were culture in DMEM/F12 plus 10% FBS as reported previously ^8,12^. Culture media were replenished every third day. Secretome from CSSC and fibroblasts were collected in log phase as reported before by culturing cells in a basal medium devoid of serum and any growth factors ^18,54^. Harvested secretomes from both CSSC and fibroblasts were further concentrated to 25X via centrifugation at 5,000 rpm in Amicon Ultra 100k cutoff ultrafilters. Concentrated secretome was mixed in a 1:1 ratio with 10mg/ml fibrinogen and applied to 0.5µl of 100U/ml thrombin to form fibrin gel on the mouse corneas.

### Mouse corneal injury and secretome treatment

All experimental procedures were reviewed and approved by the University of Pittsburgh Institutional Animal Care and Use Committee and animals were handled according to guidelines provided in the Association for Research in Vision and Ophthalmology Resolution on the Use of Animals in Ophthalmic and Vision Research. We estimated the sample size based on the variability of different assays. Sample size for mice cohorts was estimated by power analysis before initiation of the study. Both male and female mice were included in the study and randomly assigned to control and treatment groups. The experiment included four groups: 1) uninjured naïve normal controls; 2) mice received wound and fibroblast secretome; 3) mice received wound and CSSC secretome; and 4) mice received wound and sham (fibrinogen and thrombin to form fibrin gel without secretome). Twenty mice in each group were anesthetized by intraperitoneal injection of ketamine (50 mg/kg) and xylazine (5 mg/kg) and only one eye of each mouse was treated. One drop of proparacaine hydrochloride (0.5%) was added before debridement for topical anesthesia. A trephine was used to mark the central 2 mm of the cornea. An AlgerBrush II was passed over the area marked by the trephine to perform corneal epithelial debridement. After the removal of the epithelium, the AlgerBrush was applied again to damage the basement membrane and the anterior stromal tissue as described previously ^19^. Immediately after wound and sham or secretome treatment, mice received ketoprofen (3 mg/kg) for analgesia. 0.5 μl of thrombin (100 U/ml) was added to the wounded cornea, followed by 1 μl of 1:1 secretome and fibrinogen (10mg/ml) mixture or fibrinogen only as sham control. After 1 minute, a second round of thrombin and fibrinogen was applied to the cornea. One drop of gentamicin ophthalmic solution (0.3%) was added to prevent bacterial infection.

### Cell viability staining

Cells were stained with two viability dyes, Calcein Red-Orange and Hoechst 33342 (Invitrogen) after secretome harvesting from both CSSC and fibroblasts to assess cell viability. Both cell types were washed once with phosphate buffered saline (PBS) and incubated for 15 minutes in dark with the Calcein Red-Orange (1:1000) and Hoechst 33342 (1:2000). Staining was captured using 361nm/565nm excitation/emission wavelengths under an inverted fluorescent microscope (TE 200-E, Nikon).

### Flow Cytometry

For stem cell characterization, cells were washed briefly with PBS, fixed and permeabilized wherever required (for nuclear and cytoplasmic antibodies), blocked in 1% bovine serum albumin (BSA) for an hour, stained with following antibodies-CD90-BV510, CD73-PE/Cy7, CD105-AF647, OCT4-FITC, CD166-FITC, ABCG2-APC, NOTCH1-PE, STRO1-AF647, and CD271-AF647. Following isotype controls were used for the stem cell characterization experiment, IgG1 K Iso FITC, IgG2a K Isotype PE, IgG1 K Iso PE/CY7, IgG1 K Iso APC. At least 20,000 cells were acquired using FACS Aria (BD Biosciences) and analyzed using FlowJo_V10 software. At least three different CSSC strains from 3 different donors were characterized for stem cell marker expression. For cell death and apoptosis analysis post secretome harvesting, cells were stained with annexin V for apoptosis and 7-AAD for necrosis in annexin V binding buffer using 30 minutes incubation in dark. Cells were directly acquired after that using flow cytometer. For flow cytometry analysis of inflammatory and immune cells, three days following treatment, corneas were examined and rinsed, minced, and treated with collagenase type I (84 U/cornea) for 60 min at 37°C, and triturated until no apparent tissue fragments remained. The single-cell suspension of each cornea was then filtered through a 40-μm cell strainer cap (BD Labware, Bedford, MA) and washed. Corneal cells were treated with anti-mouse CD16/CD32 (Fc III/II receptor; 2.4G2) to prevent nonspecific antibody binding and then stained for various leukocyte surface markers for 30 min at 4°C. The following antibodies were used: CD45-PerCP, CD11b-AF700, GR1-PE, F4/80-PECy7, together with the Violet Live/Dead dye. The cells were analyzed via a Cytoflex cytometer and further analysis was carried out using FlowJo software. Gates were set based on staining with the appropriate single antibody and a mixture lacking that particular antibody, fluorescence minus one (FMO) control.

### Optical Coherence Tomography (OCT) scanning and analysis

Mouse corneas were evaluated using optical coherence tomography (OCT) to examine the epithelium regeneration 3 days after the treatments. A modified Bioptigen spectral-domain OCT system (Bioptigen Inc., Durham, NC, and SuperLum Ltd., Ireland) was used to acquire image volumes of mouse corneas in vivo. Scans sampled a 3 by 3 by 3 mm region of tissue with 512 by 180 by 1,024 measurements. 12-20 eyes for each group were examined by determining quadrant-scans along four axes (0°–180°, 45°–225°, 90°–270°, and 135°–315°) to ensure scanning through the central cornea and data along the 0° to 180° axis was used for analysis. The OCT images were then evaluated via FIJI (NIH). For OCT analysis mice were grouped according to their epithelial healing: 1) healed (the cornea is comparable to that of mice with no wounding); 2) partially healed (slight opaqueness of the cornea/area outside of pupil not completely healed), or 3) not healed (the cornea is with ulcer/too thin/too thick). The data was then analyzed via chi-squared analysis at the alpha = 0.05 level.

### Immunofluorescence and Whole Mount Staining

For immunofluorescence staining, fixed corneas 12µm cryosection were used. 6-8 corneal sections on each slide were permeabilized using 0.1% triton-X for 15 minutes and blocked using 1% BSA for 1 hour. Cornea sections were incubated in primary antibodies overnight and secondary antibodies for 2 hours at room temperature using 4′,6-diamidino-2-phenylindole (DAPI) as nuclear staining. The primary antibodies used for the staining of cornea sections are as follows-CD45-eF450, F4/80-APC, GR1-PE, CD59-FITC, collagen IV. SPARC, α-SMA, collagen 3A1, vitronectin. After staining, slides were acquired using laser scanning confocal microscope and analyzed using FV10-ASW4.2 Viewer. Corneas were acquired from both central and peripheral positions and fluorescence intensity was averaged to give final mean fluorescence intensity (MFI) value using Image J.

For wholemount staining mouse corneas were fixed for 1 hour in 1.3% paraformaldehyde in PBS at room temperature. Radial incisions were made to allow for flat mounting of the corneal tissues. Corneas were washed in PBS five times, permeabilized in 1% Triton-X-100 in PBS at room temperature for 60 minutes and blocked with 20% goat serum in blocking buffer (0.3% Triton-X-100/0.1% Tween-20 in PBS) for 1 hour. The corneas were then incubated in a mixture of primary antibodies β-3 tubulin and P-substance for 2 hours at room temperature, followed by an additional incubation overnight at 4°C. After five 5-minute washes in a buffer (0.1% Tween-20 in PBS), the corneas were incubated in a mixture of following secondary antibodies in a blocking buffer at room temperature for 2 hours. Following five 10-minute washes with a wash buffer, the corneas were mounted on slides and dried at 4°C for at least 12 hours before imaging. Images were acquired in multiple z-stacks by sequential scanning on FV1200 confocal microscope and stitched on FV10 viewer. The following secondary antibodies were used for immunofluorescence and wholemount staining-Donkey anti-goat IgG AF-555, Donkey anti-rabbit AF-555 IgG, Donkey anti-mouse AF-647 IgG, Donkey anti-Rabbit AF-488 IgG, and Donkey anti-rat-488 IgG. For IgG autoantibody staining, Goat anti-mouse IgG-647 was used. The immunofluorescent experiments were performed at least in three different corneas per group and repeated 2-3 times. Mean fluorescence intensity quantifications were done by two independent observers.

### TUNEL staining

Corneal cell death detection using TUNEL staining was performed using a commercially available kit, as per manufacturer’s instructions on cryosections of corneal tissue. Nuclei were stained with DAPI. At least three independent eyes from each condition and eight sections of each condition were stained and imaged using a confocal microscope. For quantification, TUNEL+ cells were counted manually by two independent observers.

Blinded methods were used for study outcome and designs. In particular, the microscopy and histological evaluations of cornea section immunofluorescence or whole mount staining for different antibodies was performed by two independent researchers. Flow cytometry was performed and analyzed by two independent researchers. OCT was performed and analyzed by a researcher in blinded fashion. The animal experiments were repeated at least twice and secretome from at least two different CSSC and fibroblasts were used for the experiment.

### Multidimensional protein identification technology (MudPIT) analysis

Proteomic identification and analysis of the secretome was performed for both CSSC and fibroblasts. Secretome was harvested and concentrated as described above. The secretome proteins were precipitated using methanol chloroform method of Wessel and Flugge ^55^. These proteins were processed through various steps of denaturation, reduction, alkylation and digestion as reported before ^53^. Briefly, peptides obtained after the process were identified using LTQ orbitrap Elite mass spectrometer equipped with a nano-LC electrospray ionization source (ThermoScientific). Tandem mass (MS/MS) spectra were interpreted using ProluCID v. 1.3.3. DTASelect v 1.9 ^56^ and swallow v. 0.0.1 (https://github.com/tzw-wen/kite), an in-house developed software, were used to filter ProLuCID search results at estimated false discovery rates (FDRs) at the spectrum, peptide, and protein levels. All reported FDRs are less than 5%. All 4 data sets were contrasted using Contrast v 1.9 against their merged data set, using our in house developed software sandmartin v 0.0.1. ^57^. Another in-house developed software, NSAF7 v 0.0.1, was used to generate spectral count-based label free quantitation results.

**GO enrichment analysis was performed using a fee online tool**, DAVID (version DAVID 6.8; http://david.ncifcrf.gov/), and functional enrichment of the genes whose corresponding proteins have positively distributed normalized spectral abundance factor (dNSAF) were analyzed using GOstats (version 2.48.0) in both replicates of CSSC and fibroblasts. The top 10 most significantly enriched GO categories for a given cell type were compared.

R package pheatmap (version 1.0.12) was used for the **hierarchical clustering analysis and heatmap** plotting. Heatmaps were used to compare the protein expression in secretome of CSSC and fibroblasts using dNSAF protein values. In the heatmap, the row represents the name of the protein-associated gene, and columns indicate the cell type used. Similar elements were classified in groups in a binary tree using hierarchical clustering.

### Statistical analysis

All data reported in the study is presented as mean ± SD. Statistical differences were determined using one-way analysis of variance to assess the significance of differences between all groups and reported as significant p values for multiple comparisons and adjusted by the Tukey method. Few markedly deviating data points were omitted during analysis due to huge variation from rest of the samples in the group. Statistical significance was set at p < 0.05.

## Supporting information

Supplemental Figures in one pdf file

## Acknowledgments

We thank all donors and their families for donating corneas and the clinicians at UPMC Eye Center Drs. Deepinder Dhaliwal, Vishal Jhanji, Alex Mammen, and the fellows and residents who supplied the corneal rims, and the Center for Organ Recovery & Education (CORE) in Pittsburgh who supplied the corneas. We thank Nancy Zurowski for Flow Cytometry. We also thank Anthony St. Ledger for his expert suggestions on the complement study.

## Notes

http://proteomecentral.proteomexchange.org/cgi/GetDataset?ID=PXD022081

### Competing Interest Statement

The University of Pittsburgh has submitted an invention disclosure related to this study with Yiqin Du and Ajay Kumar as inventors. Other authors have declared no competing interests.

