## Supplemental Figures in one pdf file for "Stem Cell Secretome Promotes Scarless Corneal Wound Healing and Rescues Corneal Sensory Nerves"

**This PDF file includes:**

Figures S1 to S4

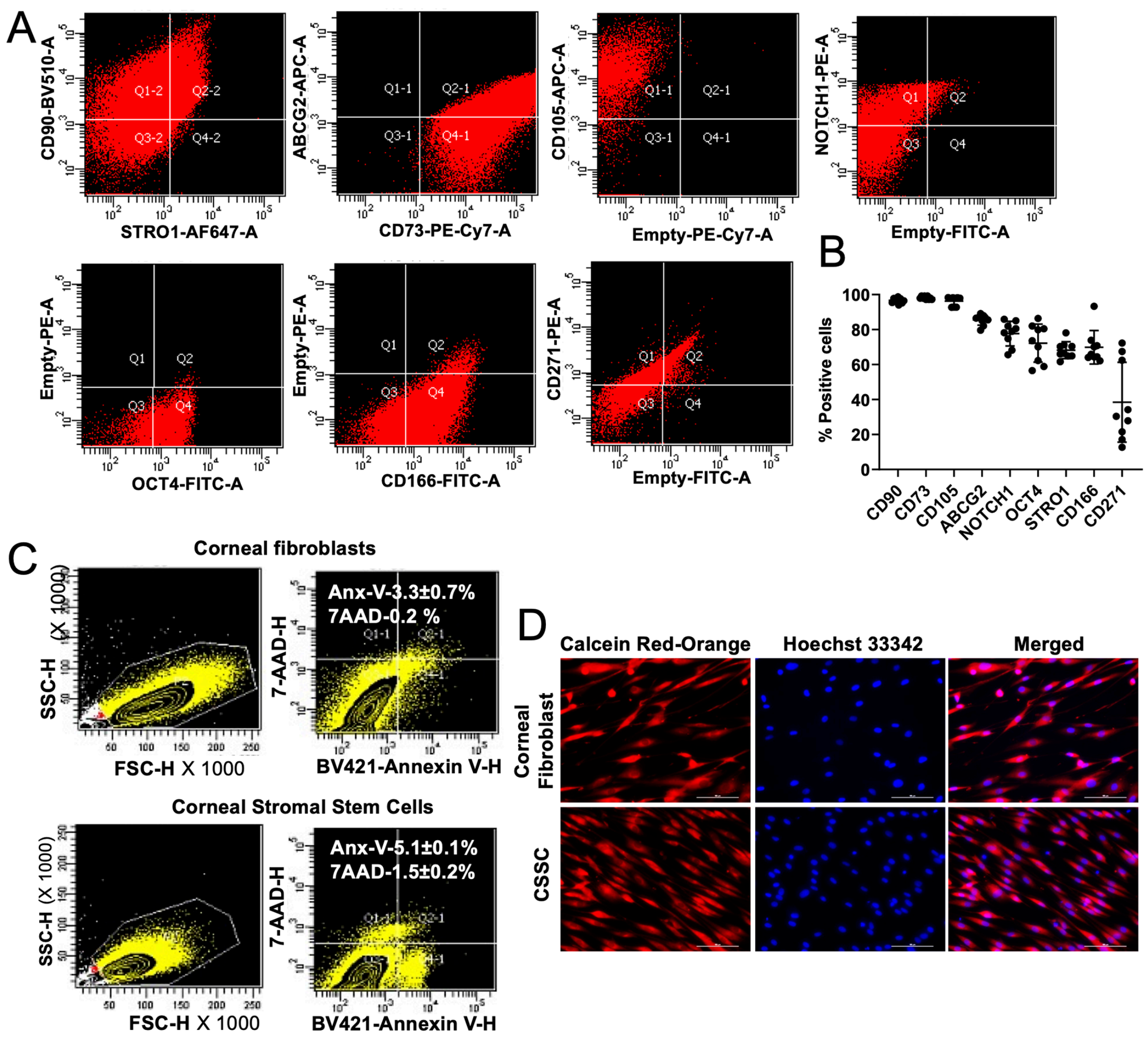


**Fig. S1. Evaluating stem cell characteristics of human corneal stromal stem cells (CSSC) and cell viability post secretome harvesting. A-B.** Dot plots from flow cytometry analysis and bar graph showing expression of stem cell markers in CSSC (n=3), **C.** Annexin V-7AAD flow cytometry analysis for cell viability assessment in corneal fibroblasts and CSSC post secretome harvesting, after incubation in serum free media. Gate was set on unstained control cells for normalizing the background fluorescence and isotype controls for elimination of non-specific staining. 100,000 events per tube were acquired. Results are representative of Mean±SD, **D.** Live cell fluorescence laser scanning confocal microscopy for cell viability in corneal fibroblasts and CSSC post secretome harvesting. Calcein Red-Orange and Hoechst 33342 were used at a dilution rate of 1:1000 and 1:2000 respectively. Scale bars-100μm.

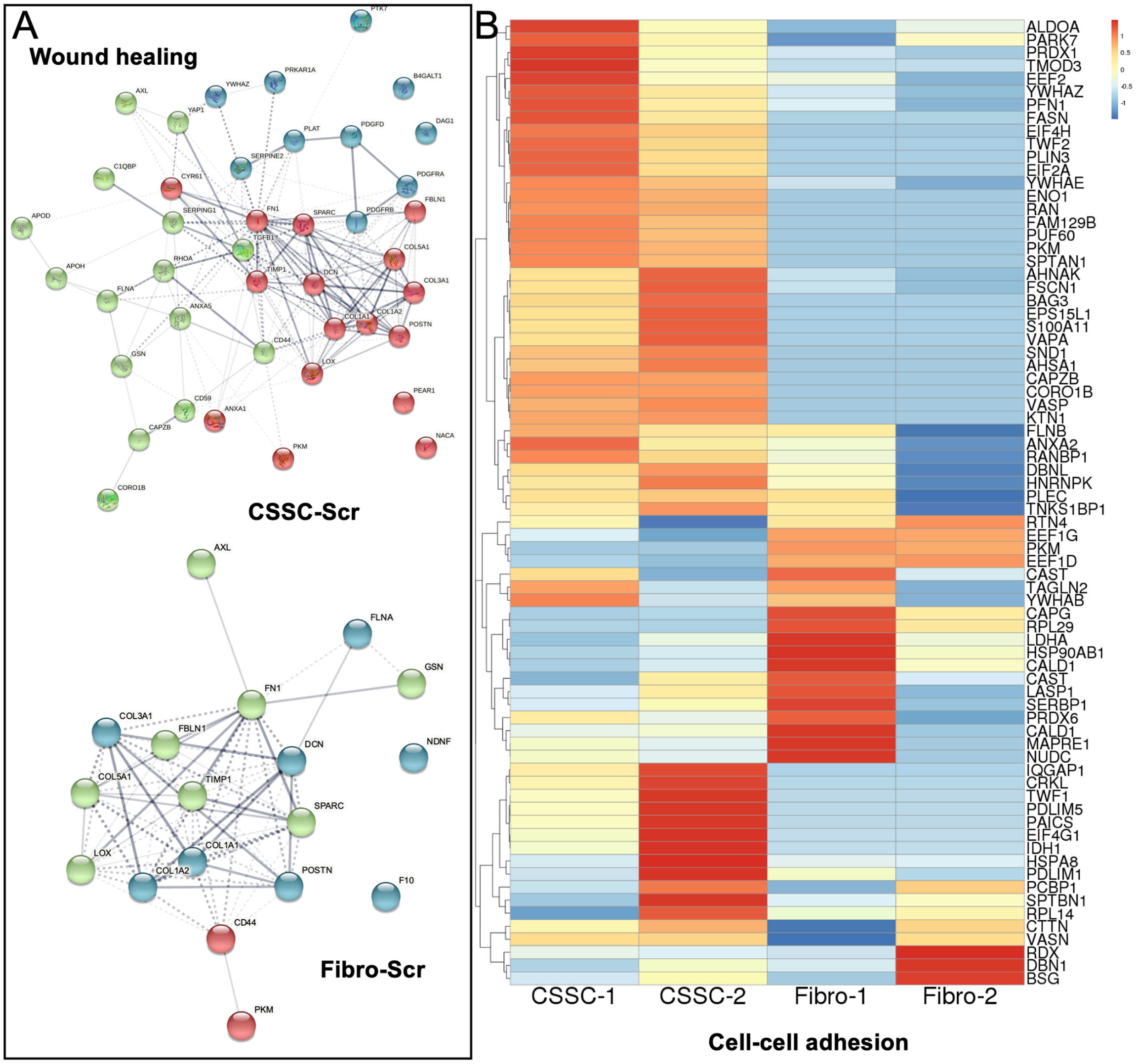


**Fig. S2. Proteomic analysis of corneal stromal stem cell and fibroblast secretomes. A.** Interactome analysis of secretome proteins from both CSSC and fibroblasts, showing interacting proteins for wound healing, more in CSSC and fewer in corneal fibroblasts, as analyzed by String v11, (n=2 each). **B.** Heatmap showing hierarchical clustering and the differential expression of cell-adhesion proteins between CSSC and corneal fibroblast secretome. Heatmaps were generated using R package pheatmap (version 1.0.12).

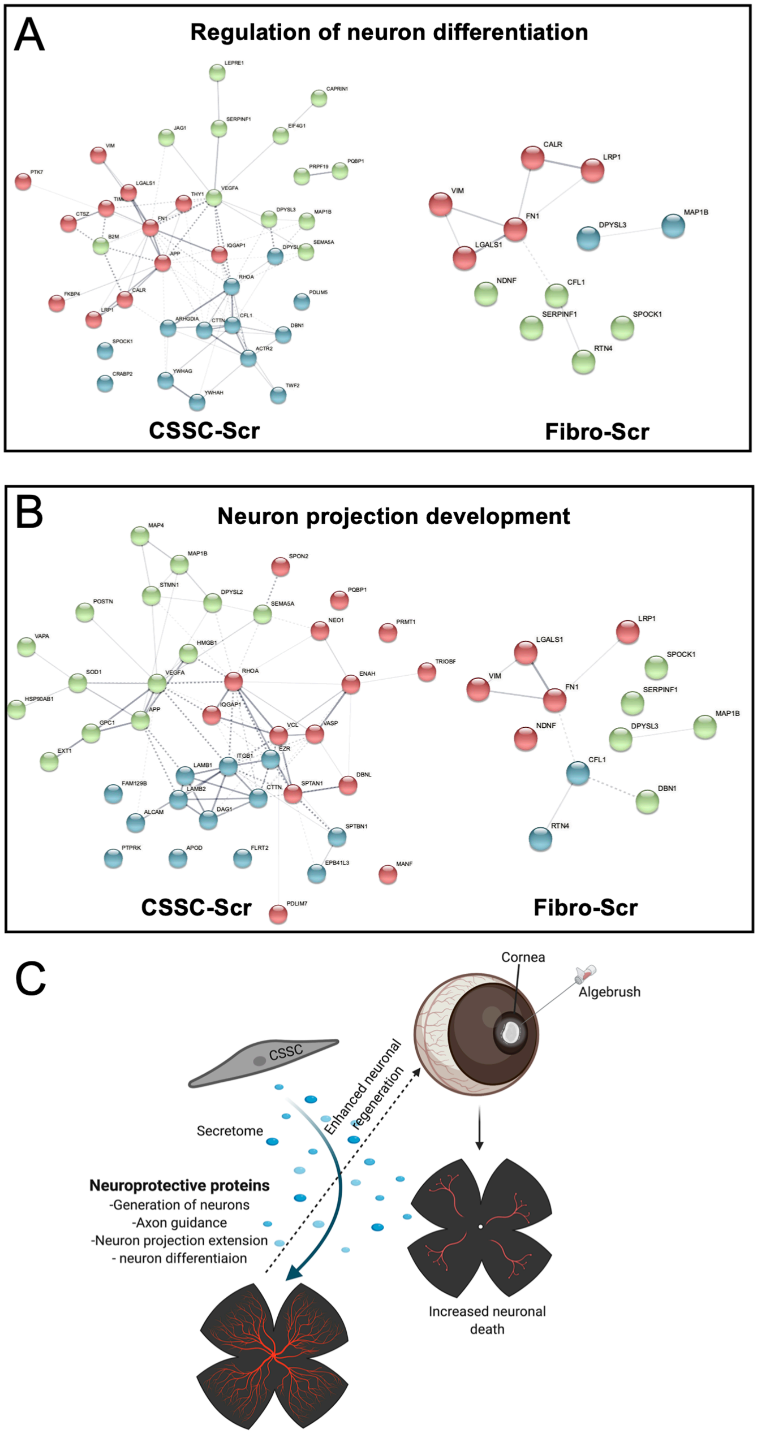


**Fig. S3. Proteomic characterization of secretome proteins involved in neuron differentiation and projection development. A.** Interactome analysis of secretome proteins from CSSC and corneal fibroblasts, showing higher number of proteins involved in regulation of neuron differentiation as compared to corneal fibroblasts, **B.** Interactome showing a similar pattern for proteins involved in the development of neuron projections, as analyzed by String v11, (n=2 each), **C.** Schematic figure showing an overall scheme for neuroprotective effect of CSSC secretome on corneal wound healing and sensory nerve regeneration (adapted from Biorender.com).

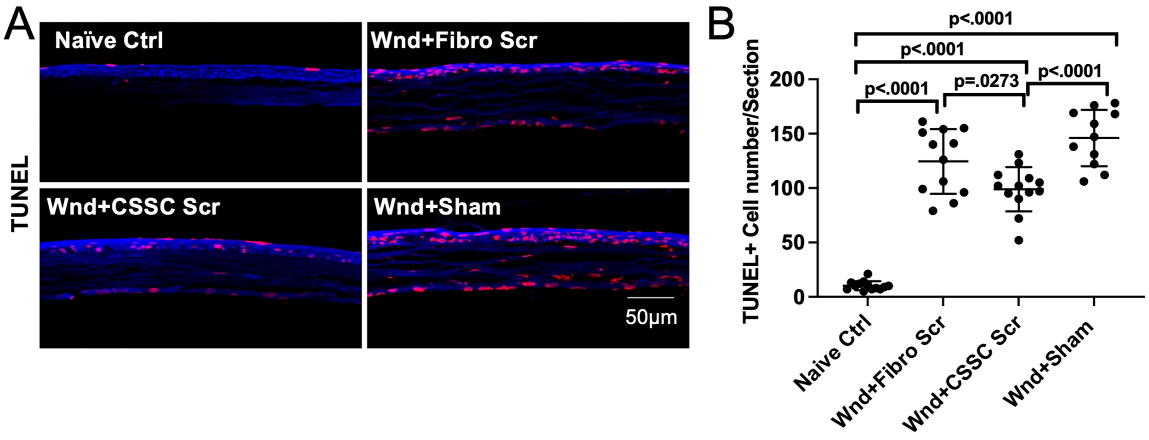


**Fig. S4. Corneal cell death analysis using TUNEL staining. A.** Immunofluorescent images showing TUNEL+ cells in the cryosections of central and peripheral corneas. The higher number of TUNEL+ cells are clearly visible in sham and fibroblast secretome treated corneas, scale bars-50μm, **B.** Dot plots showing quantification of mean TUNEL+ cells in the cornea. Data is represented as Mean±SD. Each dot on graph represents counts from one corneal section. 3-4 sections were photographed per eye (n=3 corneas).
